# Intracranial fluid dynamics changes in idiopathic intracranial hypertension: pre and post therapy

**DOI:** 10.1101/245894

**Authors:** N. Agarwal, C. Contarino, L. Bertazzi, N. Limbucci, E. F. Toro

## Abstract

Idiopathic intracranial hypertension (IIH) is a condition of unknown etiology frequently associated with dural sinus stenosis. There is emerging evidence that venous sinus stenting is an effective treatment. We use phase contrast cine MRI to observe changes in flow dynamics of multiple intracranial fluids and their response to different treatments in a patient with IIH. We quantified the following parameters at the level of the aqueduct of Sylvius and the cervical C2C3: cerebrospinal fluid (CSF), arterial and venous flow; CSF velocity amplitude; artero-venous delay (AVD); artero-CSF delay and percentage of venous outflow normalized to total arterial inflow (tIJV/tA). Analyses were run before lumbar puncture (LP) (A), after LP (B), after medical therapy (C) and after venous stent placements deployed at two separate times (D and E). AVD and tIJV/tA improved only after CSF removal and after stent placements. CSF velocity amplitude remained elevated. Arterial flow profile showed a dramatic reduction after LP with improvement in mean venous flow. This report is the first to demonstrate interactive changes in intracranial fluid dynamics that occur before and after different therapeutic interventions in IIH. We discuss how increased intracranial venous blood could be “tumoral” in IIH and facilitating its outflow could be therapeutic.

## Introduction

Idiopathic intracranial hypertension (IIH) is a neurological condition of unknown etiology, which requires prompt diagnosis and if left untreated could result in rapid progressive visual loss (Mollan *et al.*, 2016). A high opening cerebrospinal fluid (CSF) pressure during lumbar puncture (LP) is indicative of increased intracranial pressure (ICP). Increased CSF opening pressure coupled with normal neuroimaging findings are generally sufficient to make a definite diagnosis of IIH in obese females of childbearing age (Friedman *et al.*, 2013). Periodic removal of CSF through LP or the use of CSF diversion procedures are widely accepted therapeutic interventions to reduce ICP in these patients. Adjunct medical therapy consists of the use of carbonic anhydrase inhibitors or topiramate and shows little success.

Intracranial volume is maintained constant (Monroe-Kellie) in a rigid skull encasement, during each cardiac cycle (CC), through a complex temporal synchronization of multicompartment intra and extracranial fluid dynamics (Beggs *et al.*, 2013). How increased CSF pressure in the subarachnoid space (SAS) influences intracranial arterial and venous fluid dynamics within the framework of the Monroe-Kellie hypothesis remains unclear. Bateman showed that in IIH, the total cerebral arterial inflow is increased by 21% and the correspondent percentage of venous outflow through superior sagittal sinus (SSS) is reduced (Bateman, 2008b). On the venous front, there is evidence that almost 93% of patients with IIH harbor some degree of dural sinus stenosis (Morris *et al.*, 2017). Stenosis can be intrinsic or extrinsic, dynamic or static. While still debatable, there is growing evidence that CSF removal during LP causes transient resolution of dural venous sinus stenosis, suggesting that these are more likely extrinsic (Buell *et al.*, 2017; Juhász *et al.*, 2017; Rohr *et al.*, 2007). The increased resistance to venous outflow will further augment CSF pressure in the SAS by obstructing its absorption at the level of arachnoid granulations, thus leading to a vicious cycle. Improvement of venous outflow by use of stents can facilitate venous outflow and CSF absorption (Aguilar-Pérez *et al.*, 2017; R. I. Farb *et al.*, 2003).

Phase contrast cine MR (PCC-MR) is a consolidated non-invasive technique often employed in clinical settings to allow quantification of CSF flow parameters in the aqueduct of Sylvius (AoS) and determine therapeutic strategies (Stoquart-El Sankari *et al.*, 2008; Yousef *et al.*, 2016). It can provide useful information regarding velocity, flow, amplitude and direction of CSF flow but also in the arteries and veins (Enzmann and Pelc, 1993; Lagana *et al.*, 2014).

In this study, we set to observe changes in the temporal coupling and dynamics of arterial, venous and CSF compartments using PCC-MR in a patient with IIH. We repeated these analyses at different time points. The results were compared with data acquired on a healthy subject and with reported values in the literature.

## Methods

A 34 yr old obese female reported multiple episodes of neck pain, throbbing headache and progressive vision loss. LP was performed in a seated position and the opening CSF pressure was read using a hand held manometer attached to the needle. Visual acuity was assessed through Snellen charts. Visual field (VF) pattern was studied by standard automated perimetry and retinal fiber nerve layer (RFNL) thickness was measured using optical coherence tomography (OCT). Standard venous digital angiography was performed and venous pressure was measured at various locations by attaching a pressure transducer to the microcatheter.

### MRI acquisition

secondary causes were ruled out using standard diagnostic sequences (T1-SE, T2-TSE, FLAIR, DWI) using a 1.5 Tesla MR scanner (Siemens, Erlangen, Germany). Gadolinium-based venography (MRV) was performed to study venous sinus anatomy. Flow study was performed using a retrospective pulse-gated gradient echo sequence with a TR/TE of 22.9/7.0 ms. The 2D single 5mm slice of the PC sequence was positioned orthogonal to the direction of flow at the AoS (aCSF) and in the anterior CSF space between the C2 and the C3 vertebral level (cCSF) with a VENC of 20/50cm/s (CSF/vessels). 40 phases were acquired corresponding to one CC. Cardiac triggering was performed prospectively with finger plethysmography. The same protocol was run before LP (A), after 7 hours of LP (B) and after two months of standard medical treatment (C). The protocol was repeated after one month of venous stent placement in the transverse sinus (TS) (D) and again after one month of stent placement in the right TS (E). All acquisitions were performed by the same neuroradiologist. The same protocol was collected on a healthy 26 yr old male.

### Four fluid compartment analysis

CSF and blood flow quantifications were performed using SPIN (Signal Processing in NMR, Detroit, MI). Post-processing was done using MATLAB (The MathWorks, Natick, MA). Flow and velocity profiles were smoothed using built-in functions of MATLAB. A region of interest (ROI) was manually drawn for each vessel in the magnitude images of the PCC. Over the CC, the size, shape and location of each ROI were kept fixed. At C2C3 level we manually drew 7 ROIs: left/right internal jugular veins (IJVs), internal carotid arteries (ICAs), vertebral arteries (VAs) and one additional ROI for the phase offset (supplementary Fig. 1). For CSF (aCSF and cCSF), we manually drew a ROI on the phase image corresponding to the highest caudal velocity and one additional ROI for the phase offset. The cross-sectional area of the ROI was assumed to remain constant across all time points. Velocity amplitude of the aCSF and the cCSF was calculated from the maximum and minimum peaks of their absolute values. Arteriovenous delay (AVD) was estimated as the lag in time between arterial and venous systolic peaks and was represented as a percentage of CC. Artero-CSF delays were estimated as the lag in time between arterial systolic peak and aCSF (AaCD) and cCSF (AcCD). To assess the contribution of collateral pathways for venous blood flow exiting the intracranial compartment, we calculated the ratio, expressed in percentage, between the total internal jugular vein outflow (tIJV) and the total intracranial arterial inflow (tA), that is tIJV/tA=tIJV/(ICAs+VAs) (Lagana *et al.*, 2014). When tIJV/tA is close to 100%, venous blood outflow through the IJVs matches the arterial blood inflow, and therefore the contribution of collaterals is minimum. On the contrary, when tIJV/tA decreases to zero, major contribution is given by collateral pathways (Doepp *et al.*, 2004).

## Results

### I) Pretreatment data (A)

#### Clinical status

neurological exam was normal. Opening LP pressure was 50 cmH_2_0 (normal value in the seated position is <34.6 cmH_2_0) (Abualenain **et al.**, 2011) that dropped to 24cmH_2_0 after removing 30mL of CSF. *Ophthalmologic findings:* visual acuity was reduced in both eyes (right eye 20/100, left eye 20/25). Severe bilateral papilledema was noted with slit-lamp examination. *MRI and MRV:* MRI was negative for space occupying disease. Enlarged optic nerve sheaths were found bilaterally: 7.2mm on the right side and 8.4mm on the left side (normal values are 5.2 ± 0.9 mm) (Passi *et al.*, 2013). 3D MRV showed a left TS stenosis. *Angiographic procedure:* bilateral stenosis at the transverse-sigmoid sinus junction was observed with pressure difference of 4 mm Hg across the left TS and only 1 mmHg across the right TS. Only the left TS was stented by placing a 7×50 mm Carotid Wallstent that lowered the difference to 1 mmHg (D) (Supplementary Fig. 2).

#### PCC-MRI

The temporal order of all four systolic peak flows was conserved in the patient with respect to both our healthy subject and that reported in the literature (Fig. 1a) (Beggs, 2014). The AVD was 15.35% of CC and in the normal range (Table 1). Likewise, the AaCD and AcCD were also in the normal range (20.74% and 5.35% respectively) (Table 2). aCSF velocity amplitude in the patient was almost three times higher than that of healthy subject (12.24cm/s vs. 4.15cm/s) (Supplementary figure 3). cCSF caudal peak velocity and flow were within the normal range (−2.93cm/s and −2.96mL/s respectively) (Table 2). While mean arterial flow was in the normal range (13.35mL/s), the systolic peak showed a longer plateau (Fig. 2A). Mean venous flow, in absolute value, was below normal minimum values (5.88mL/s) (Table 1). The tIJV/tA was 44% with respect to 93.5% in the healthy subject.

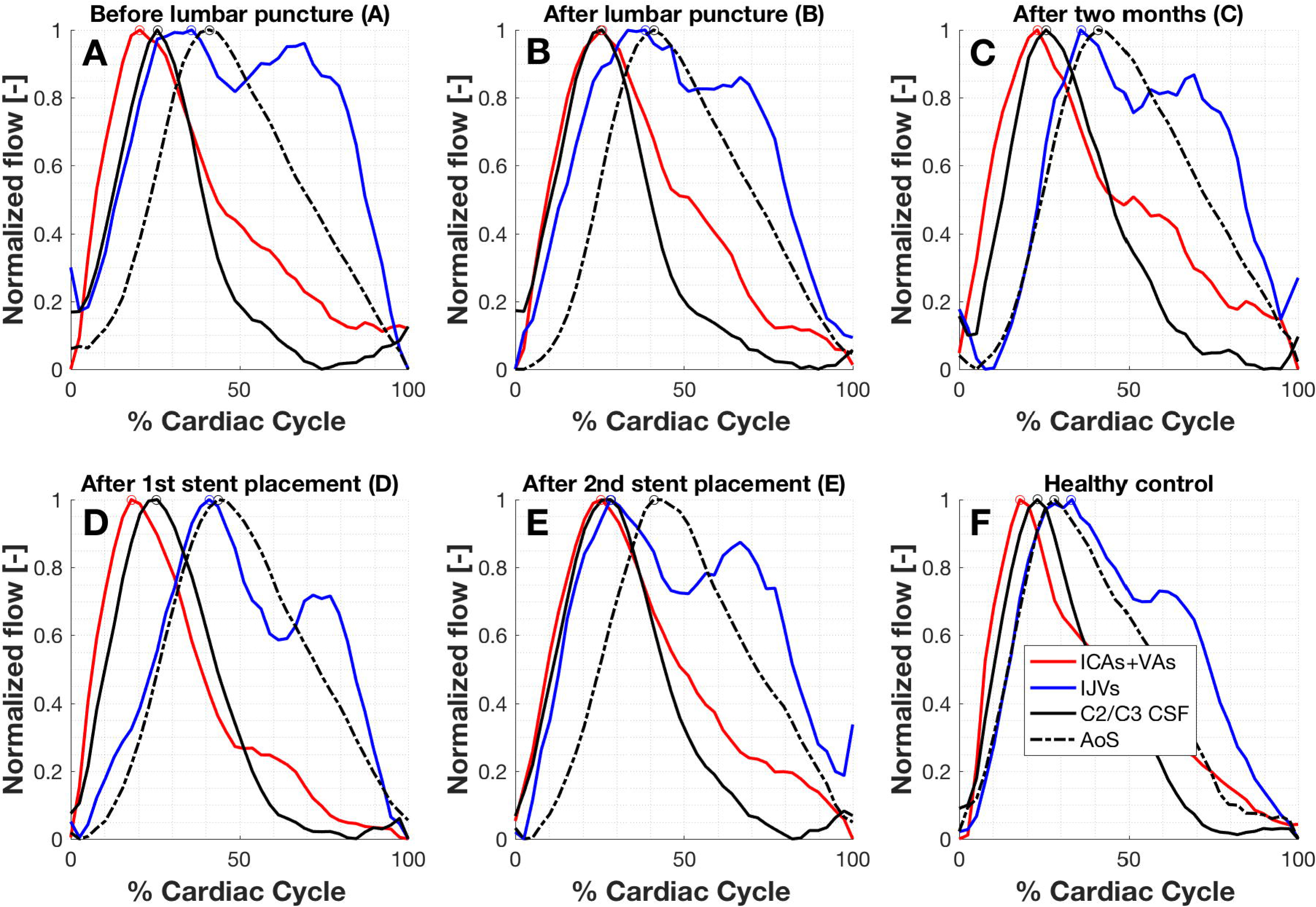

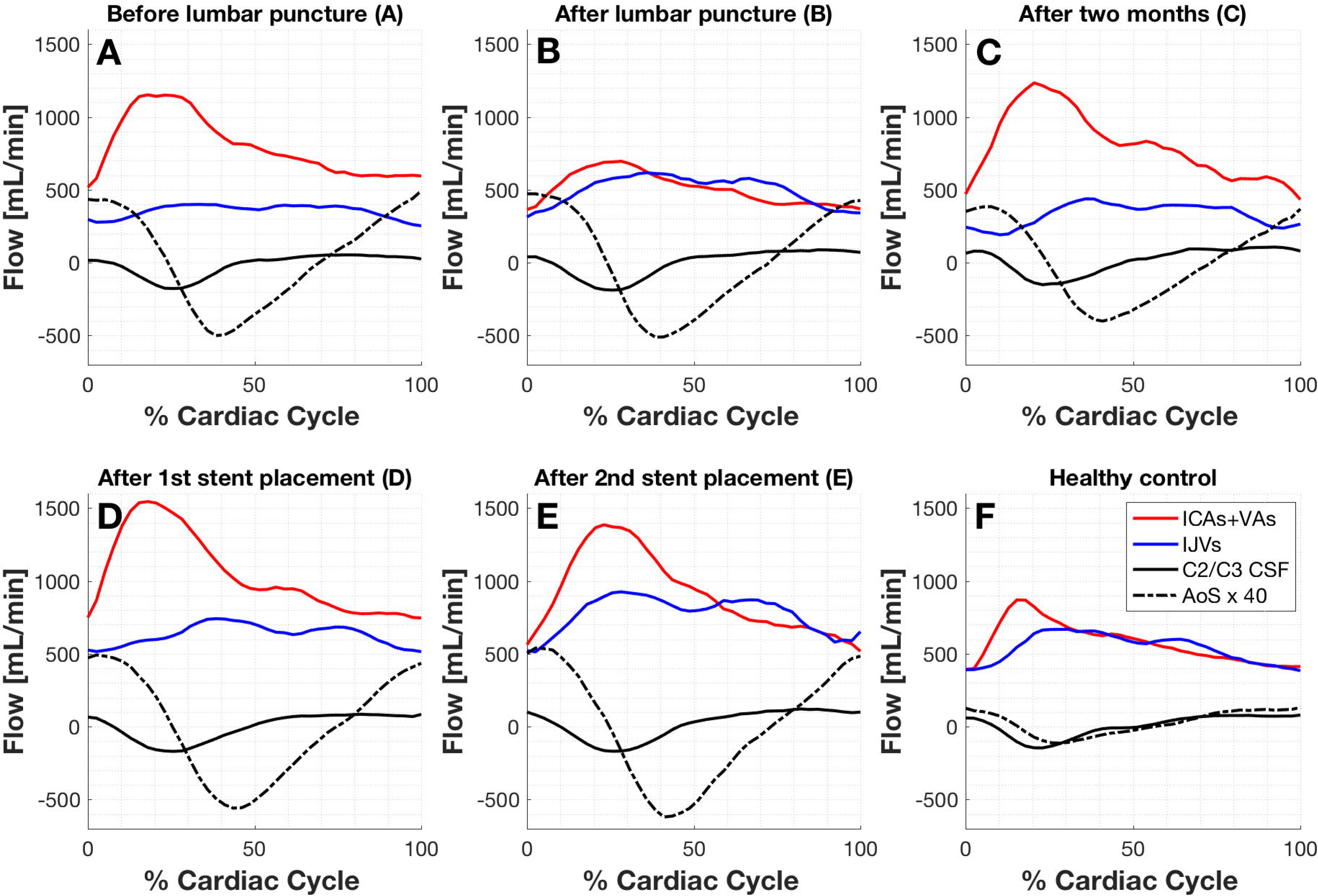

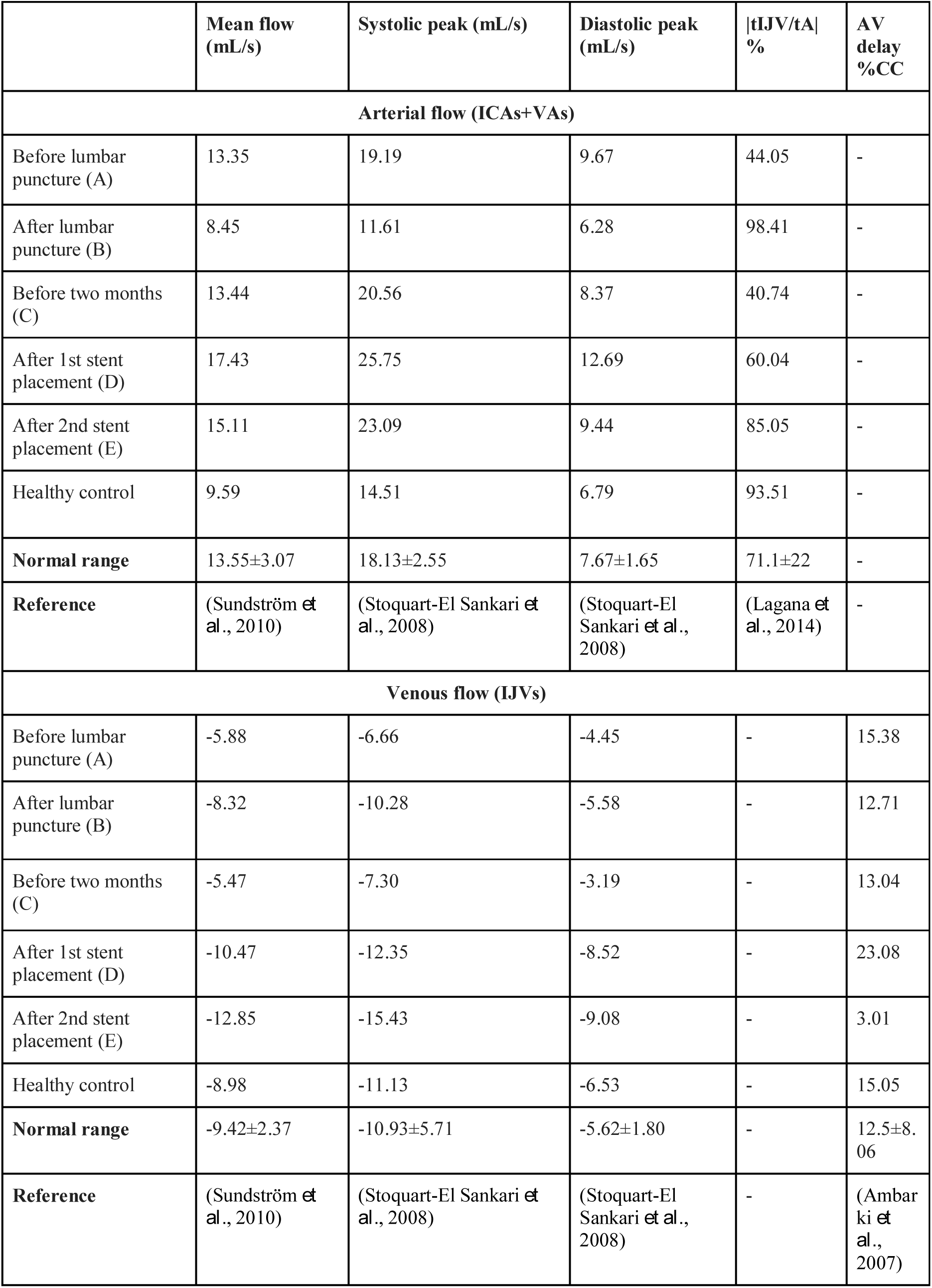

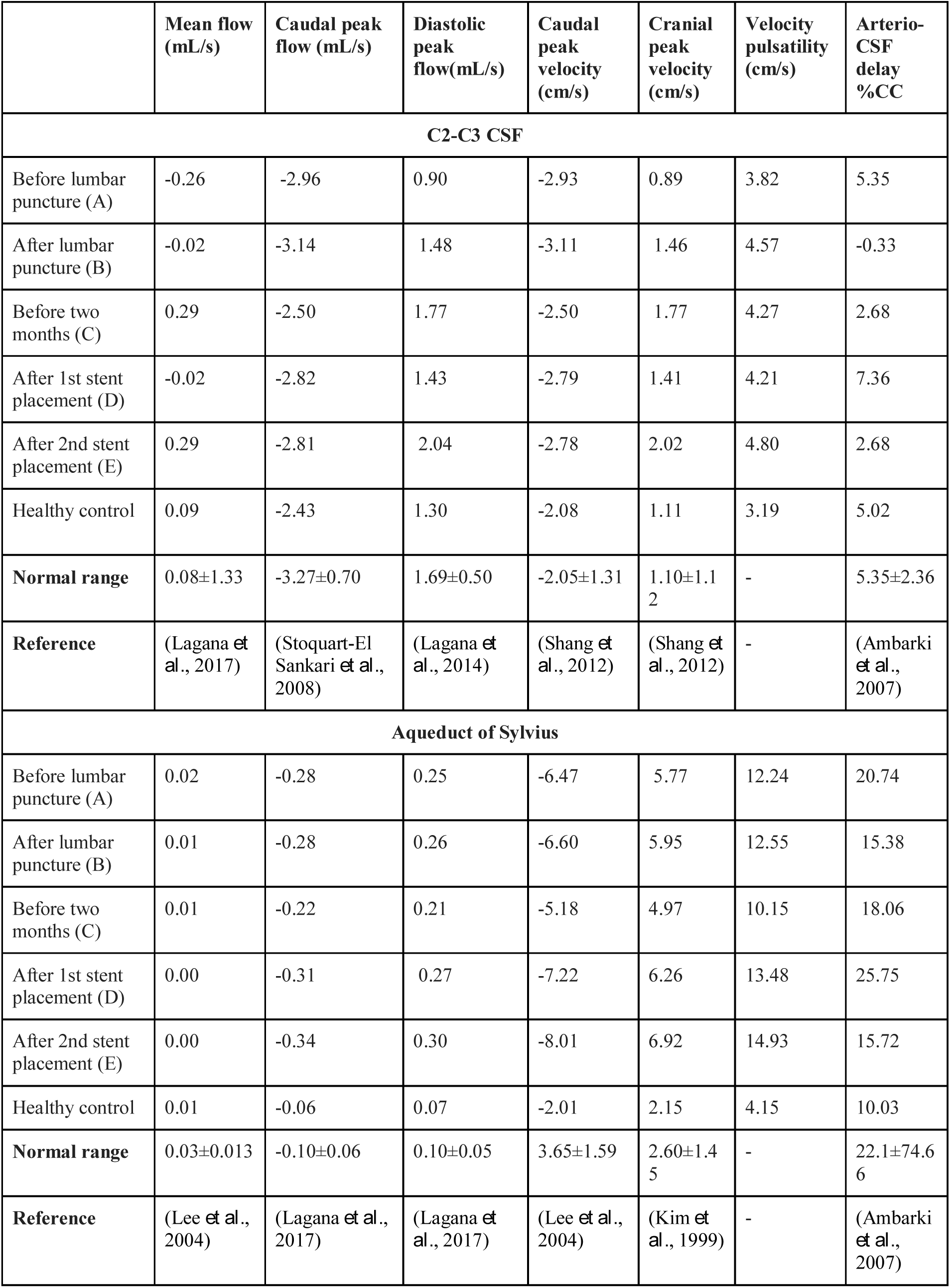

|  | Mean flow<br>(mL/s) | Caudal peak<br>flow (mL/s) | Diastolic<br>peak<br>flow(mL/s) | Caudal<br>peak<br>velocity<br>(cm/s) | Cranial<br>peak<br>velocity<br>(cm/s) | Velocity<br>pulsatility<br>(cm/s) | Arterio-CSF<br>delay<br>%CC |
| --- | --- | --- | --- | --- | --- | --- | --- |
| **C2-C3 CSF** |  |  |  |  |  |  |  |
| Before lumbar<br>puncture (A) | -0.26 | -2.96 | 0.90 | -2.93 | 0.89 | 3.82 | 5.35 |
| After lumbar<br>puncture (B) | -0.02 | -3.14 | 1.48 | -3.11 | 1.46 | 4.57 | -0.33 |
| Before two<br>months (C) | 0.29 | -2.50 | 1.77 | -2.50 | 1.77 | 4.27 | 2.68 |
| After 1st stent<br>placement (D) | -0.02 | -2.82 | 1.43 | -2.79 | 1.41 | 4.21 | 7.36 |
| After 2nd stent<br>placement (E) | 0.29 | -2.81 | 2.04 | -2.78 | 2.02 | 4.80 | 2.68 |
| Healthy control | 0.09 | -2.43 | 1.30 | -2.08 | 1.11 | 3.19 | 5.02 |
| **Normal range** | 0.08±1.33 | -3.27±0.70 | 1.69±0.50 | -2.05±1.31 | 1.10±1.1<br>2 | - | 5.35±2.36 |
| **Reference** | (Lagana *et al.*, 2017) | (Stoquart-El Sankari *et al.*, 2008) | (Lagana *et al.*, 2014) | (Shang *et al.*, 2012) | (Shang *et al.*, 2012) | - | (Ambarki *et al.*, 2007) |
| **Aqueduct of Sylvius** |  |  |  |  |  |  |  |
| Before lumbar<br>puncture (A) | 0.02 | -0.28 | 0.25 | -6.47 | 5.77 | 12.24 | 20.74 |
| After lumbar<br>puncture (B) | 0.01 | -0.28 | 0.26 | -6.60 | 5.95 | 12.55 | 15.38 |
| Before two<br>months (C) | 0.01 | -0.22 | 0.21 | -5.18 | 4.97 | 10.15 | 18.06 |
| After 1st stent<br>placement (D) | 0.00 | -0.31 | 0.27 | -7.22 | 6.26 | 13.48 | 25.75 |
| After 2nd stent<br>placement (E) | 0.00 | -0.34 | 0.30 | -8.01 | 6.92 | 14.93 | 15.72 |
| Healthy control | 0.01 | -0.06 | 0.07 | -2.01 | 2.15 | 4.15 | 10.03 |
| **Normal range** | 0.03±0.013 | -0.10±0.06 | 0.10±0.05 | 3.65±1.59 | 2.60±1.4<br>5 | - | 22.1±74.6<br>6 |
| **Reference** | (Lee *et al.*, 2004) | (Lagana *et al.*, 2017) | (Lagana *et al.*, 2017) | (Lee *et al.*, 2004) | (Kim *et al.*, 1999) | - | (Ambarki *et al.*, 2007) |

### II) Post-treatment data (B, C, D and E)

#### Clinical status

no clinical improvement was noticed by the patient at B and at C. Mild relief in headache was referred at time point D but immediate benefit was reported after time point E. *Ophthalmologic findings:* there was a trend towards progressive decrease in visual function at time points B and C whereas at time points D and E all visual function including RNFL parameters appeared stable in both eyes with slight visual improvement in the left eye (data not shown). *MRI and MRV:* no morphologic changes were observed at any time point except a 1 mm reduction in the left optic nerve sheath diameter at E. MRV did not change at B and C. At D, the right TS was severely reduced in diameter. *Angiographic procedure:* two months after D, a new right TS stenosis with a pressure difference of 6 mm Hg difference was found and a second stent was deployed (E) (Supplementary fig. 2).

#### PCC-MRI

AVD reduced slightly at B (12.71%), without significant change at C (13.04%) (Fig. 1b). There was a dramatic increase in AVD at D (23.08%) that reduced greatly after the second stent placement, E (3.01%) (Table 1). aCSF velocity amplitude in the patient remained high at all time points (table 2; supplementary fig. 2). Similarly, cCSF caudal peak velocity and flow remained constant all time points (table 2; supplementary fig. 2). Arterial flow showed a marked transient reduction after LP (8.45mL/s) (Fig. 2B). It returned to normal range values (13.55 ± 3.07mL/s) at C, D and E (13.44mL/s, 17.43mL/s and 15.11mL/s) (Table 1). Mean venous flow normalized at B (−8.32mL/s) but reduced to −5.47mL/s at C, improving again after stent placements to −10.47mL/s and −12.85mL/s. tIJV/tA improved markedly after LP to 98.41%, albeit transiently, since at C the percentage outflow reduced back to 40.74%. This percentage improved at D and E to reach 60.04% and 85.05% respectively (Table 1).

## Discussion

To our knowledge, this is the first study that describes simultaneous interactive changes in multiple fluid compartments of the brain and spine in a patient with IIH before and after medical and interventional therapies using a validated PCC-MR technique.

During systole, arterial pulsations in pial arterioles are transmitted directly into the incompressible CSF filled SAS. This transmitted pulse evokes a serious of events in the following temporal order: a) CSF in SAS shifts out of the foramen magnum; b) venous blood from the sinuses egresses out of the brain and c) part of the CSF from the ventricles is displaced out towards the spinal SAS (Ambarki *et al.*, 2007). During the diastolic phase, venous outflow decreases and venous blood is stored in the highly compliant thin-walled cortical and bridging veins that drain into the SSS (Bateman, 2003). An increase in SSS pressure would hamper venous outflow from cortical and bridging veins. In addition, there will be increased collateral flow and increased mean transit time, that would redirect the volume of venous blood through a longer path within the cranial cavity (Mancini *et al.*, 2012). It is reasonable to suggest that this might result in a “permanent” increased volume of venous blood in the cranial cavity. Could this extra “stored” blood in the cortical veins be “tumoral”?

By applying Darcy’s basic law of flow we can relate flow q (volume transferred per unit time) to pressure difference Δ*p* = *p_u_* − *p_d_* and resistance *R*, namely

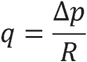

Here *p_u_* denotes upstream pressure and *p_d_* denotes downstream pressure. We note that this law ignores the pulsatile, time dependent nature of flow and relies solely on averaged values. Bateman invoked this law to relate ICP to CSF outflow *q_CSF_*, from the cerebral SAS into the superior sagittal sinus (SSS), and to pressure *p_SSS_* in SSS as follows:

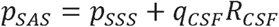

with *R_CSF_* denoting resistance to CSF outflow (Bateman, 2008a). Invoking again Darcy’s law of flow, we can express the total cerebral venous outflow *q_CVO_* from the brain into the right atrium as

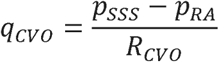

where *p_RA_* is the pressure in the right atrium and *R_CVO_* is the resistance to venous flow from the brain to the heart. We note that this is a very rough approximation to a very complex process but nonetheless gives some useful insight. By combining the last two equations and noting that ICP is accepted to be *p_SAS_* we can express ICP as

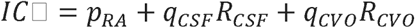

In the event of an increase in the external pressure (ICP or *p_SAS_*, assuming frozen values for the remaining values, the sinus wall will collapse and increase *R_CVO_* and vice versa. Increased *R_CVO_* can be reduced either indirectly by removing CSF through LP or CSF diversion procedures or directly by placing a venous sinus stent that will mechanically keep the vessels patent, facilitating venous outflow. The added benefit in the second case is an improvement in CSF reabsorption into SSS in addition to being a long term effect (Aguilar-Pérez *et al.*, 2017)

The most important complication, although rare, is remote or stent-adjacent stenosis (Raper *et al.*, 2017). Our patient developed a high-pressure difference in the right TS suggesting the presence of continued increased external CSF pressure. This underscores the importance of considering the entire cerebral venous system to avoid collapse of previously normal compliant veins (Bateman, 2008b).

PCC-MRI flow analysis in our patient provided us useful additional information regarding changes in fluid dynamics. Despite increased ICP, our results show that the overall fluid dynamics and temporal order of all fluid systolic peaks were normal before treatment (Fig. 1a), although AVD, AaCD and AcCD were slightly prolonged. We deduce that all fluid compartments respond physiologically, albeit with longer time delays. Medical treatment alone did not change these time delays, but both CSF removal and stent placements greatly enhanced the response of fluids to arterial input (Fig. 1). In addition, and most importantly, blood flow analysis shows that both CSF removal and venous stenting greatly improves mean venous outflow. This data suggests that our patient had extrinsic reversible stenosis.

The dramatic reduction in the arterial flow profile after LP is curious and we do not have an explanation for this observation. However we conjecture that there might be a degree of arterial hyperemia in line with Bateman’s observation regarding increased arterial inflow in IIH (Bateman, 2008b). By reducing *R_CVO_* by CSF removal, a possible new setting of pressure changes and flow dynamics through cerebral autoregulatory mechanisms might in part explain such changes (Miller *et al.*, 1972; Moriyasu and Hayashi, 1978).

The limitations of this study are linked to data derived from a single patient study. However, all variables in our patient were kept constant to monitor changes making comparisons easy. Since all interventions were intended for treatment, some results at later time points might reflect cumulative effects of previous treatments.

## Conclusions

A thorough analysis of interactive intracranial fluid dynamics may provide the key in understanding what causes increased ICP in IIH and how best to treat it. Increased resistance to cerebral venous outflow may result in “storage” of extra intracranial venous volume. Venous stenting may be an efficacious strategy to improve egression of cerebral venous blood in the long-term.

## Acknowledgements

The authors thank Prof. E. M. Haacke for providing us the software, Signal Processing in NMR, for CSF and flow analysis.

## Abbreviations

AaCD: artero-aqueduct CSF delay
AcCD: artero-cervical CSF delay
aCSF: CSF through the AoS
AoS: aqueduct of Sylvius
AVD: arterovenous delay
CC: cardiac cycle
cCSF: CSF in the cervical subarachnoid space
CSF: cerebrospinal fluid
CVO: cerebral venous outflow
ICA: internal carotid artery
ICP: intracranial pressure
IIH: idiopathic intracranial hypertension
IJV: internal jugular vein
tA: total cerebral arterial inflow
tIJV: total outflow from internal jugular veins
LP: lumbar puncture
RFNL: retinal fiber nerve layer
PCC-MR: phase contrast cine magnetic resonance
SAS: subarachnoid space
TS: transverse sinus
VA: vertebral artery

